# A novel CLAVATA1 mutation causes multilocularity in *Brassica rapa*

**DOI:** 10.1101/2022.09.28.509982

**Authors:** Hiu Tung Chow, Timmy Kendall, Rebecca A. Mosher

**Affiliations:** School of Plant Sciences, The University of Arizona, Tucson, AZ 85721, USA

**Keywords:** Brassica rapa, CLAVATA-WUSCHEL, locule, seed set, floral meristem, fruit, gynoecium

## Abstract

Locules are the seed-bearing structure of fruits. Multiple locules are associated with increased fruit size and seed set, and therefore control of locule number is an important agronomic trait. Locule number is controlled in part by the CLAVATA-WUSCHEL pathway. Disruption of either the CLAVATA1 receptor-like kinase or its ligand CLAVATA3 can cause larger floral meristems and an increased number of locules. In an EMS mutagenized population of *Brassica rapa*, we identified a mutant allele that raises the number of locules from 4 to a range of from 6 to 8. Linkage mapping and genetic analysis support that the mutant phenotype is due to a missense mutation in a *CLAVATA 1* (*CLV1*) homolog. In addition to increased locule number, *brclv1* individuals fail to terminate their floral meristems, resulting in internal gynoecia that negatively impact seed production.

## Introduction

In angiosperm fruit, locules are the chamber surrounding the pericarp that contains the seeds. Locules are derived from the fusion of carpels, the innermost whorls of a flower and the female reproductive structure (reviewed in Herrera-Ubaldo and de Folter, 2022). During Brassicaceae flower development, carpels fuse into a cylindrical gynoecium separated by thin septa that form locules. Cells in the margins between fused carpels give rise to the ovules, resulting in a row of ovules in each locule. Gynoecia can form from the fusion of two or more carpels, resulting in two or more carpel margins. After fertilization, the gynoecium elongates and differentiates into a silique (the fruit), and the walls of each carpel are known as valves. Generally, the number of valves reflects the number of carpels originally present in a gynoecium, and the number of locules is often equivalent to or slightly fewer than number of valves.

The early stage of floral development requires the *CLAVATA* (*CLV*) signaling pathway (Somssich *et al*., 2016). In *Arabidopsis thaliana* (Arabidopsis), *CLV1* encodes a Leucine-rich repeat (LRR) receptor-like kinase perceiving a small peptide ligand encoded by *CLV*3 (Trotochaud *et al*., 2000; Clark *et al*., 1997). Mutations disrupting *CLV1* cause a range of abnormalities in floral tissues including extra carpels and ectopic floral organs (Diévart *et al*., 2003; Clark *et al*., 1997). *clv1* mutants also display a range of silique phenotypes including club-shaped siliques, partial valves/valveless, and extra valves. Mutation of *CLV3* causes multicarpel gynoecia and siliques with extra valves (Fiers *et al*., 2006; Song *et al*., 2013). The abnormalities observed in *clv1* and *clv3* mutants are attributed to the disruption of stem cell maintenance, which results in enlarged floral meristems, which results in more cells contributing to carpel development, as well as increased cell division in the valve margins (Durbak and Tax, 2011).

Because ovules arise from the carpel margins, ovule/seed number is correlated with carpel/locule number (Herrera-Ubaldo and de Folter, 2022; Xu *et al*., 2021). The more carpels in the gynoecium, the more cells have the meristematic properties to allow ovule and seed formation. This hypothesis has been tested mostly in the *Brassica* genus, where 2-carpel gynoecia and bilocular siliques are the ancestral state, but some varieties have additional locules. A naturally-occurring single nucleotide polymorphism (SNP) in *Brassica rapa* (*B. rapa*) *CLV3* (resulting in a Pro-to-Ler substitution at amino acid 582) causes tetralocular siliques and higher seed set (Yadava *et al*., 2014; Fan *et al*., 2014). Similarly, multilocularity in a *Brassica juncea* cultivar is controlled by natural variation in 2 *CLV1* homoeologs (Xiao *et al*., 2013; Xiao *et al*., 2018; Chen *et al*., 2018). Induced mutation of the two *CLV3* homoeologs in *Brassica napus* results in multilocular siliques with higher seed yield and higher 100-seed weight than either single mutant or wild type. However, unlike the natural variants, these induced loss-of-function mutations have additional pleiotropic effects (Yang *et al*., 2018).

Here we show that a single missense mutation in A07p048430.1_BraROA (*BrCLV1)* increases locule number in *B. rapa*. However, in contrast to the prevailing hypothesis, the multilocular fruits of this mutant do not result in greater seed production due to additional internal gynoecia that adversely affect seed set. This research provides alternative insights into the engineering of multilocularity to optimize seed yield.

## Results

### Isolation of a novel multilocular mutation in *B. rapa*

While most varieties of *B. rapa* have the ancestral bilocular character, R-o-18 is tetralocular due to a non-synonymous substitution in *CLV3* (Katiyar *et al*., 1998; Fan *et al*., 2014) (**Figure 1A-B**). In an EMS mutagenized R-o-18 M3 population, we found a mutant that forms additional locules, resulting in siliques that are wider than R-o-18 (**Figure 1C-D**). We named this mutant *pomona* after the Roman goddess of fruit. Quantifying valve number in dried siliques demonstrated that *pomona* increased valve number from 4 to 6 (±0.9) (**Figure 1E**). The developing flowers on *pomona* individuals also had extra petals, anthers, and carpels, with enlarged gynoecia and fasciated stigmas (**Figure 1F-H**). Cross-sections revealed that ectopic gynoecia began to form inside the primary gynoecium in all individuals. Approximately 16% of flowers displayed more severe abnormalities, including cracked and twisted gynoecia resulted from incompletely fused carpels, and multiple internal gynoecia (**Figure 1I**). The growth of additional gynoecia suggests that the carpel meristem failed to terminate appropriately, which is commonly seen in *clv* mutants especially *CLV3* alleles (Sablowski, 2007).

**Figure 1.**
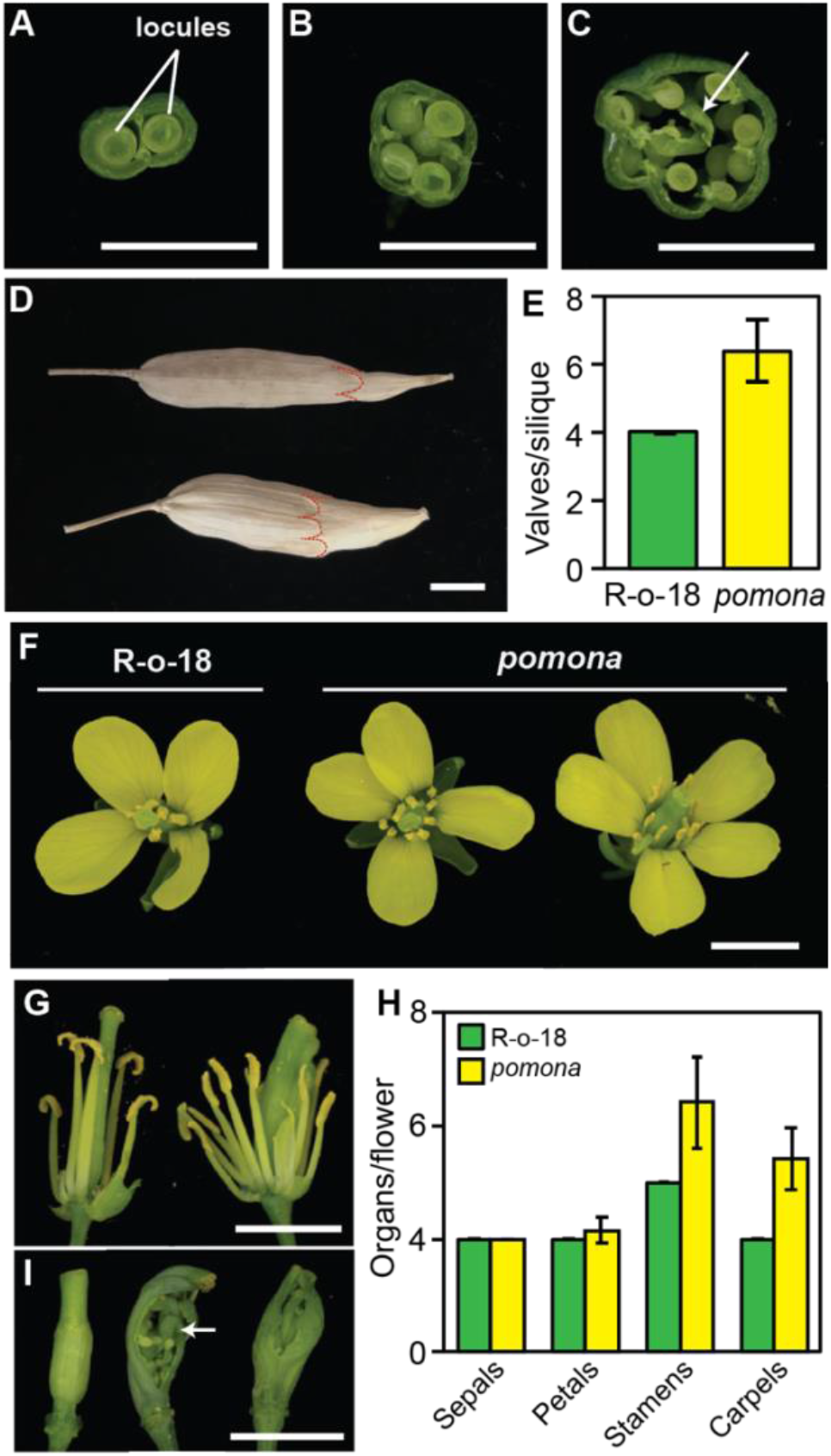
*pomona* displays multilocular phenotype. (A) Simplified floral structure in *Brassica rapa*. (A-C) Cross-section of stage 18 developing siliques from the bilocular variety R500 (A), the tetralocular variety R-o-18 (B), and *pomona* in the R-o-18 background (C). The white arrow points to the ectopic gynoecium growing inside a *pomona* primary gynoecium. Scale bar = 0.5 cm. (D) Images of complete mature and dried siliques of R-o-18 (top) and *pomona* (bottom) before shattering. The top edges of the valves are marked with dashed red lines. Scale bar = 0.5 cm. (E) Average number of valves per silique in 180 mature dried siliques from 10 individuals in *pomona* and R-o-18. Error bars represent standard deviation. (F) Represent images of R-o-18 and *pomona* flowers. (G) Enlarged images with petals removed of gynoecia in R-o-18 (left) and *pomona* (right). (H) Average number of floral organs in R-o-18 and *pomona* flowers. Quantification of 150 flowers with enclosed gynoecia as the carpel number in the cracked gynoecia could not be accurately counted. Error bars represent standard deviation. (I) In *pomona* flowers, 84% of the gynoecia were well-enclosed (left) but 16% displayed cracked ovary walls (middle and right), with ectopic gynoecia growing inside (white arrows). Scale bar = 0.5 cm.

To further characterize the mutation, we first crossed *pomona* to R500 (a bilocular *B. rapa* variety). All F1 individuals (n=10) were bilocular, suggesting that *pomona* is a recessive mutation. Alternatively, the additional valve phenotype of *pomona* might depend on the *CLV3* allele from R-o-18 (*CLV3*^R-o-18^). To test these hypotheses, F1 plants were allowed to self-fertilize, and we conducted a linkage analysis in the F2 population. First, we scored valve number in 331 F2 individuals. The F2 population displayed 3 phenotypic classes: 221 individuals had 2 valves, 64 individuals had 4 valves, and 46 individuals had 5-10 valves (the *pomona* phenotype). We eliminated another 33 individuals for which we were unable to score for valve number due to other developmental defects. These defect might result from additional EMS-derived mutations in the *pomona* background, and might explain the lower-than-expected number of *pomona*-like individuals in the F2 population. Using a derived cleaved amplified polymorphic sequence PCR marker, *CLV3* alleles were identified in the expected Mendelian ratio (11 *CLV3*^*R500*^ homozygous, 17 heterozygous, 18 *CLV3*^*R-o-18*^ homozygous, p-value < 0.05) among the *pomona-*like individuals, indicating that the *pomona* phenotype is independent from *CLV3* genotype and is genetically unlinked from *CLV3*.

### A missense allele of a *CLV1* ortholog is a candidate for *pomona*

To map the mutation, we performed mapping-by-sequencing on a DNA library constructed from pooled DNA of the 46 class III F2 individuals (**Table S1**). Analysis of the allele frequencies of R-o-18/R500 polymorphisms narrowed the causal region to a ∼4.1-Mb interval on chromosome 7 (**Figure S1**). Further variant analysis revealed 7 non-synonymous, EMS-induced SNPs (G-to-A or C-to-T) in coding regions within the interval (**Table 1**), however only SNPs in *A07p049690*.*1_BraROA* and *A07p048430*.*1_BraROA* have 100% of reads supporting the EMS-induced variants (**Table 1**, highlighted). *A07p049690*.*1_BraROA* is a putative carboxyphosphoenolpyruvate mutase (BrCPEPM; homologous to *At1g77060*; 86% nucleotide identity, e-value: 3e-163), while *A07p048430*.*1_BraROA* is *BrCLV1* (82% nucleotide identity, e-value: 0.0). Using domain similarity with AtCLV1 to annotate LRRs and the kinase domain, we determined that the mutation is located at nucleotide 1745 (CDS) within the 20^th^ LRR, causing a serine to asparagine substitution at amino acid 582 (**Figure 2A**). This is one of two *AtCLV1* homoeologs among the three *B. rapa* subgenomes (Chen *et al*., 2022). Given the prior knowledge of *clv1* mutant phenotypes, *BrCLV1* is a strong candidate for the gene disrupted in *pomona*.

**Table 1.**
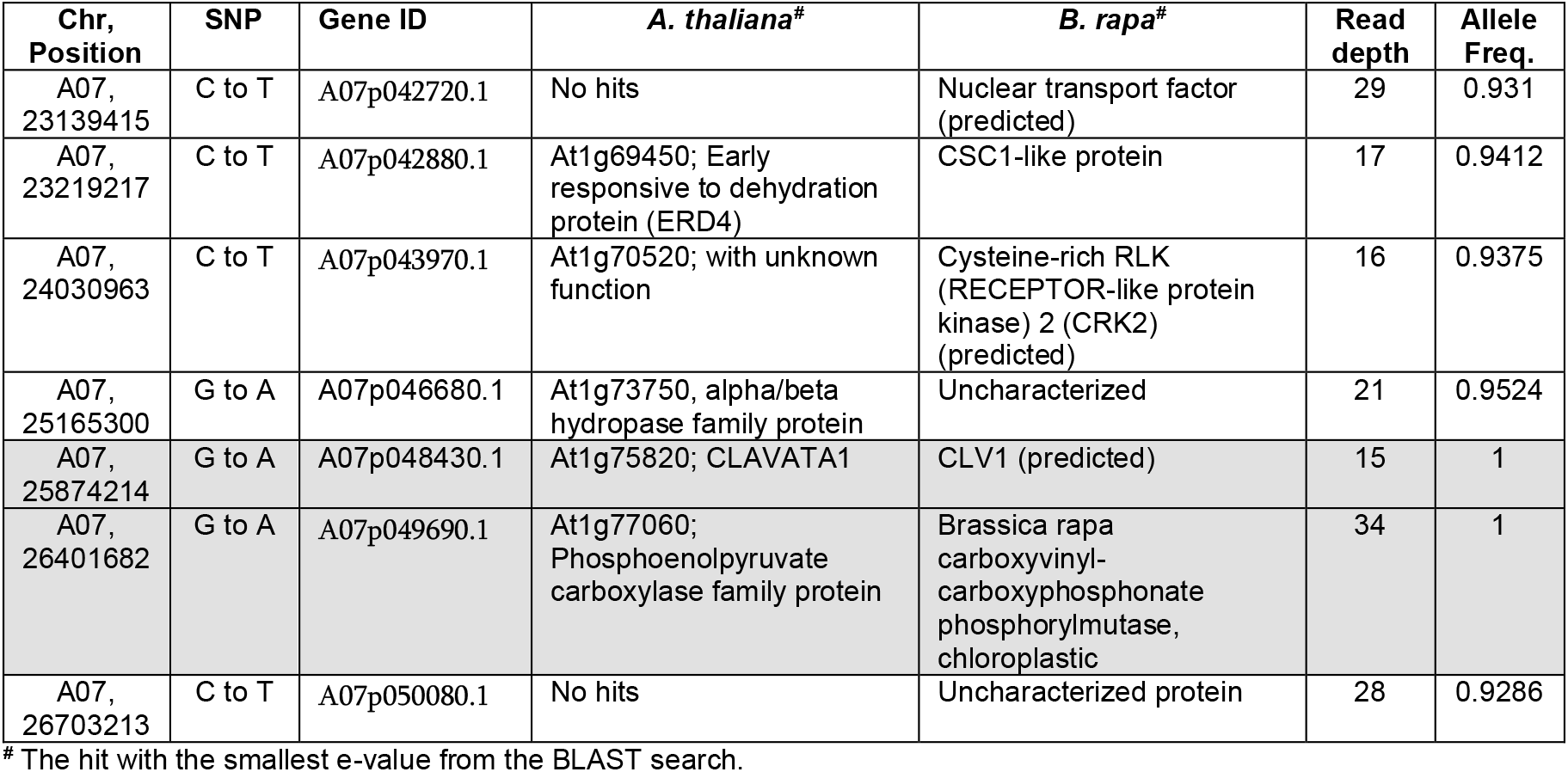
Candidates SNPs in *pomona*.

**Figure 2.**
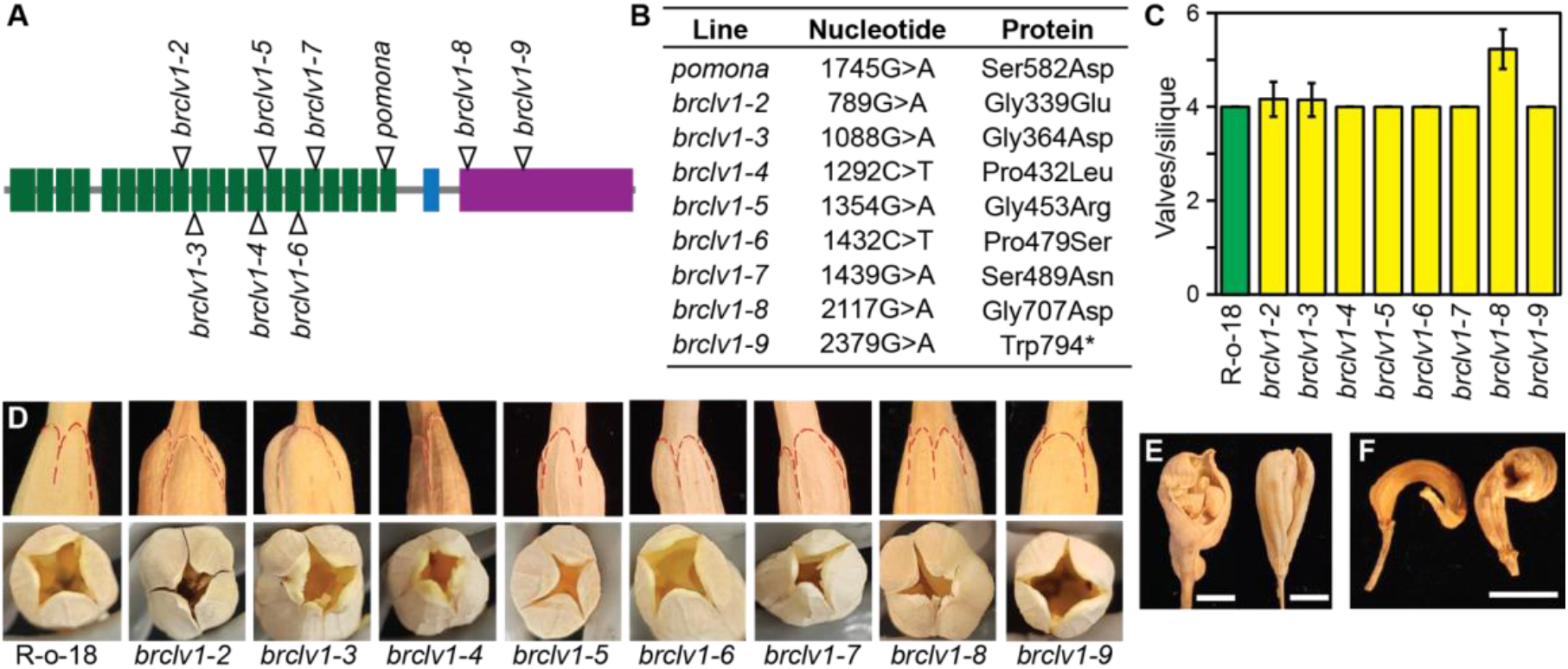
TILLING population of *A07p048430*.*1_BraROA* and allelism test. (A) Scheme of *A07p048430*.*1_BraROA*. Black, grey, and white rectangles represent LRR, intracellular, and kinase domains, respectively. Asterisk (*) represents the S582N substitution in *pomona*. Triangles indicate the verified point mutation in each allele. (B) Summary of mutation of *pomona* and the TILLING alleles. Line: allele; Nucleotide: position of the point mutation in *A07p048430*.*1_BraROA* CDS. Protein: position and the substituted amino acid, and * represents stop codon. (C) Average number of valves per silique were counted in at least 75 siliques per genotype from 5 individuals. Error bars represent standard deviation. (D) Photos are enlarged portion of siliques. To have better view of valve margins, the first row they are marked with red dashed lines, while bottom row is the aerial look of siliques which beaks were removed. In addition to completely fused siliques showing here for *brclv1-2* and *brclv1-3*, these lines often displayed cracked siliques as shown in Figure 3E. (E) Cracked siliques in *brclv1-2* (left) and *brclv1-3* (right). Scale bar = 0.5 cm. (F) The F1 of *pomona* x *brclv1-9*. Representative images of the F1.

To further investigate linkage between these two candidate mutations and the multilocular phenotype, we grew an additional 183 F2 individuals and genotyped them for *BrCLV1* and *BrCPEPM* alleles. For the BrCLV1 alleles, we observed 52, 79, and 52 homozygous mutant, heterozygous, and homozygous wild type, respectively. We then phenotyped floral development and observed that the *pomona* phenotype was restricted to homozygous *brclv1* individuals. Forty-nine of 52 *brclv1* homozygotes displayed the *pomona* phenotype; the 3 remaining had more severe developmental phenotypes and failed to produce any siliques. At *BrCPEPM* we identified 49 wild-type, 84 heterozygous, and 50 homozygous individuals. Two of the homozygous *brcpepm* mutants lacked the *pomona* phenotype and 4 *brcpepm* heterozygotes display the *pomona* phenotype. Taken together, this observation indicates that mutation of *BrCPEPM* is not responsible for the multilocular phenotype, and that the *pomona* mutation is within 0.3 cM of *BrCLV1*.

### Additional mutations in BrCLV1 influence silique phenotypes

To functionally validate that a mutation in *BrCLV1* is responsible for the multilocular phenotype, we ordered additional mutations in *A07p048430*.*1_BraROA* from a sequenced R-o-18 TILLING population (Stephenson *et al*., 2010) and bred 8 mutations to homozygosity. The alleles were renamed from *brclv1-2* to *brclv1-9* (**Figure 2A-B**). To examine evolutionary conservation at the mutated residues, we aligned BrCLV1 amino acid sequence with CLV1 orthologs in 15 eudicots (**Figure S2**). To study the effects of these mutations, we examined the phenotypes of siliques and quantified valve numbers. The *brclv1-8* allele displayed the strongest increase in the number of valve formation with (5.23 ± 0.42) (**Figure 2C-D; Figure S3**). While *brclv1-2* and *brclv1-3* alleles produce just slightly additional valves, these lines displayed shorter beaks (the apical portion of the fruit arising from the style) and club-shaped siliques, and produced additional gynoecia, frequently resulting in cracked siliques **(Figure 2E**; **Figure S3**). The remaining 5 homozygous mutations *(brclv1-4, brclv1-5, brclv1-6, brclv1-7*, and *brclv1-9*) had no observable effect on silique development or valve number (**Figure 2C-D; Figure S3**).

While most of the identified mutations are missense mutants, *brclv1-9* is a nonsense mutation at the catalytic kinase domain which creates a premature stop codon and is expected to result in a loss-of-function allele. We confirmed that neither homozygous nor heterozygous *brclv1-9* plants have any observable silique phenotype. To further confirm that the multilocular phenotype is due to mutation of *A07p048430*.*1_BraROA*, we crossed *pomona* with *brclv1-9* and examined the F1 progeny. If the *A07p048430*.*1_BraROA* is not the causal mutation, F1 progeny should be heterozygous for both *pomona* and *BrCLV1* and have wild-type siliques. While F1 individuals from this cross had siliques with four locules, they shared characteristics with *pomona*, including abnormal siliques and multiple gynoecia. Importantly, none of the F1s from this cross developed healthy siliques (**Figure 2F**). Since the *pomona x brclv1-9* F1s do not look entirely wild-type, these data suggest that the mutation responsible for multilocularity is allelic to *BrCLV1*.

Since *B. rapa* is not transformable, we moved *BrCLV1* transgenes into Arabidopsis *clv1* mutants (**Figure 3A**). We chose *clv1-11, clv1-12*, and *clv1-13* as they are null mutants with increased number of carpels/valves (Diévart *et al*., 2003), and we used both the wild-type and *pomona* alleles of *BrCLV1*. We also transformed the wild-type *AtCLV1* driven by the same promoter in parallel as a control. Both *AtCLV1* and *BrCLV1* wild-type transgenes complemented the mutant phenotype in *clv1-11, clv1-12*, and *clv1-13*, confirming that the transgenes were correctly expressed (**Figure 3B**). The transgene carrying the *pomona* allele complemented the mutant phenotype of *clv1-12* and *clv1-13*, both of which are in the WS background, but enhanced the mutant phenotype of *clv1-11*, which is in the Landsberg *erecta* (L*er*) background (**Figure 3B**). This finding is consistent with a previous observation that ecotype background modifies the *clv1* phenotype (Diévart *et al*., 2003; Durbak and Tax, 2011). Our experimental findings collectively support that the missense mutation in *BrCLV1* is the causal mutation leading to multilocularity in *B. rapa*. Hereby, *pomona* was renamed as *brclv1-1*.

**Figure 3.**
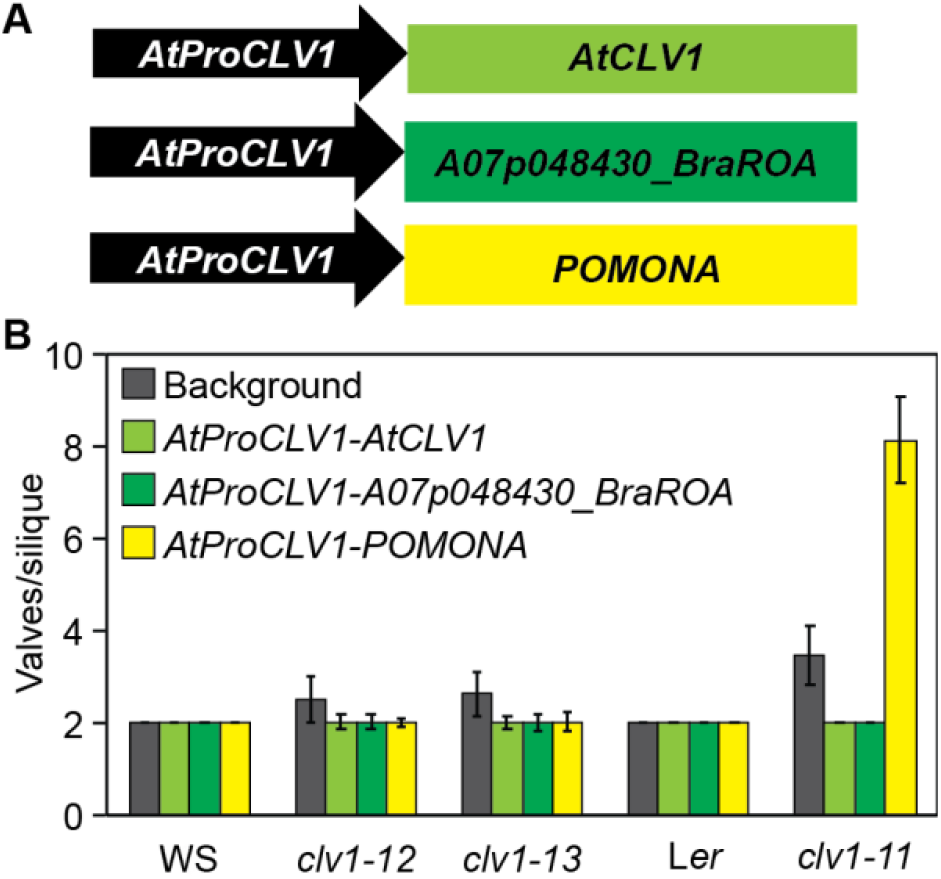
*A07p048430*.*1_BraROA* genetically complements *atclv1*. (A) Schematic diagram of the cloning constructs used for transformation. (B) Average number of valves/silique in each transformants. 15 siliques from at least 7 lines were counted in each. Columns represent each transformed construct. Error bars represent standard deviation.

### Seed yield in *brclv1-1*

The *brclv1-1* allele described here was initially identified from an M3 EMS-mutagenized population based on partial seed yield restoration in the *nrpd1a-2* background (Grover *et al*., 2018). In the mapping data, there was no significant change of allele frequency around *NRPD1a* (*A09p015000*.*1_BraROA*) on chromosome 9. *NRPD1a* also segregated as expected in the F2 mapping population, confirming that the multilocularity phenotype is independent of *NRPD1a*. Since *brclv1-1 nrpd1a-2* has higher seed yield than *nrpd1a-2* (**Figure S4**), we asked whether *brclv1-1* alone caused higher seed set. Since *NRPD1a* is required for seed development, we eliminated homozygous *nrpd1a-2* individuals in the following analyses. We then compared seed production between wild-type, heterozygous, and homozygous *brclv1-1* individuals from the F2 population. In contrast to the *nrpd1a-2* background, both seeds per silique and total seed counts demonstrate that fewer seeds were produced in *brclv1-1* homozygotes (**Figure 4A-B**). Because these individuals are from the mapping population, they are also segregating for the *CLV3*^*R-o-18*^ allele, which is associated with higher seed production (Fan *et al*., 2014). To determine the extent to which the *CLV3*^*R-o-18*^ allele modified the yield parameters in our population, we measure seeds per silique and total seed count for all *CLV3* allelic combinations. Surprisingly, *CLV3*^*R-o-18/R-o-18*^ has only subtle effects on both seed parameters in this population under these conditions (**Figure 4C-D**). Since *CLV1* and *CLV3* are in the same pathway, we further asked if *BrCLV1* acted additively with *CLV3*^R-o-18^ in controlling seed set. We tested the effect of *BrCLV1-1* alleles within the *CLV3*^*R-o-18/R-o-18*^ population. The sample size of this experiment was relatively small since 3 unlinked mutated genes were considered. In this case, there was a slight reduction (p-value < 0.05, two-sample t-test) on seed yield in individuals carrying *brclv1-1*/*brclv1-1* (**Figure S5**).

**Figure 4.**
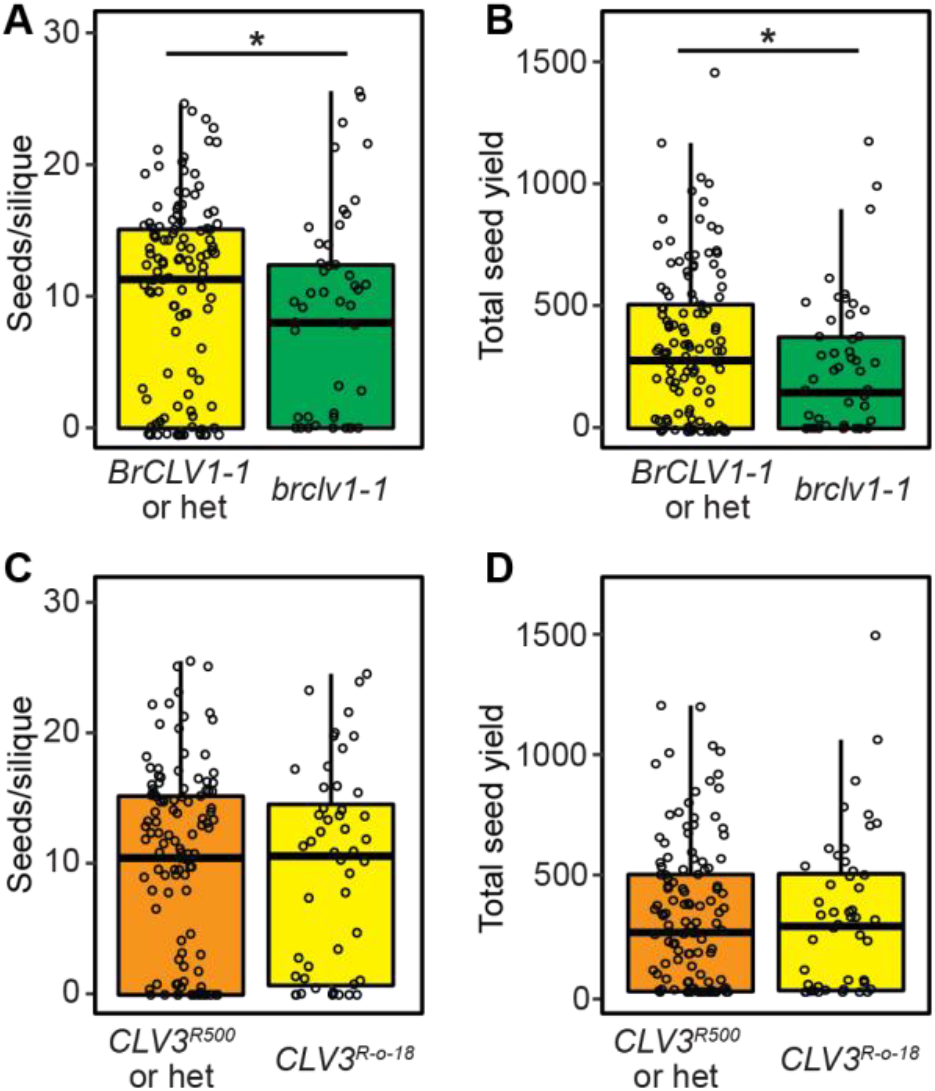
Br*CLV1* and Br*CLV3* have subtle effects in boosting seed yield. (A-B) Seed yield in Br*CLV1* alleles from the segregating F2 population. Boxplots of seeds/silique number per plant (A) and total seed yield per plant (B). Asterisk above denotes significant difference according to two-sample t-test (p-value of (A) 0.017 < 0.05; (B) 0.020 < 0.05). (C-D) Similar analysis as above but for *CLV3* alleles. No significant difference between comparing groups according to two-sample t-test.

## Discussion

An increased number of floral organs, including the carpels that develop into the locules of the fruit, can be attributed to excess accumulation of stem cells in the floral meristem or mutation of floral homeotic genes (Denay *et al*., 2017; Shang *et al*., 2019). CLV1 is involved in regulating stem cell proliferation in the shoot apical meristem (SAM). *CLV1* mutation causes an enlarged SAM and therefore more stem cells for floral meristem (FM) development, eventually causing an increased number of floral organs. Here, we demonstrate that *pomona*, a missense mutation in the *BrCLV1* LRR domain, causes increased floral organs, resulting in multilocular fruits.

The *pomona* mutation is recessive as heterozygotes are indistinguishable from wild type. However, when a *pomona* allele is in *trans* with a presumed loss-of-function allele (*brclv1-9*), the resulting plants do not completely phenocopy *pomona* homozygotes, suggesting that dosage of the mutant protein might influence phenotype in the absence of wild-type protein. Similar complexities are observed when the mutant *pomona* allele is introduced into multiple *atclv1* null backgrounds. *Brclv1-1* causes multilocularity in Arabidopsis only in the *clv1-11* background. This observation suggests that the phenotype of *brclv1-1* mutation has ecotype-specific effects, perhaps due to the *erecta* mutation in this genetic background (Diévart *et al*., 2003; Durbak and Tax, 2011). ERECTA is a LRR-receptor like kinase acting in a pathway parallel to CLV1 to confine *WUS* expression during floral meristem development (Mandel et al., 2014). Recent biochemical and genetic analysis suggests that ERECTA also serves as an upstream regulator of CLV3 in controlling shoot apical meristem maintenance (Zhang *et al*., 2021). The enhanced phenotype observed when *brclv1-1* is expressed in *clv1-11* could be due to additive ectopic WUS induction by *L*er-*ERECTA* and *brclv1*, resulting more carpels/valves formation. Our results highlight the many different phenotypes that can arise due to variation in *CLV1*.

Although multilocularity is associated with increased seed set in Brassica species, we do not detect increased seed production among the *brclv1-1* individuals tested here. One possibility is that the genetic load in these lines due to other EMS mutations could adversely reduce seed production. Such mutations might explain the wide deviation in seed production values for *BrCLV1* individuals, including many individuals that did not produce seeds (**Figure 4A**). However, we also observed that *brclv1-1* multilocular siliques consistently contained additional internal gynoecia, which occupied cavity space where the ovule/seeds should develop. These internal gynoecia, which also sometimes prevented fusion of the primary carpels, likely arose through failure to terminate stem cell activity. *CLV1* transcripts are repressed by KNUCKLES (KNU) during early floral development, resulting in termination of stem cell activity, which necessary for FM determinacy (Shang *et al*., 2021; Sun *et al*., 2019; Sun *et al*., 2009). Although the KNU pathway is not well-characterized in *B. rapa*, in Arabidopsis, KNU binds to both the promoter and first exon, a region close to the *pomona* mutation (Shang *et al*., 2021), suggesting that KNU regulation might be disrupted in *brclv1-1*.

While additional internal gynoecia and secondary EMS mutations likely underlie the reduced seed set of *brclv1-1* fruits, *brclv1-1* was originally discovered in a suppressor screen looking for increased seed production in the *nrpd1* background, indicating that *brclv1* influences seed development in some genetic contexts. The additional gynoecia occupy internal space of ovary which constraints seed set, but such restriction might be eased in the *nrpd1* mutant, which has many small aborted seeds. In addition to its well-known role in stem cell maintenance, CLV1 also localizes to plasmodesmata in the root and is hypothesized to regulate the movement of developmental factors there (Stahl *et al*., 2013; Stahl and Simon, 2013). Intercellular movement of siRNAs produced by NRPD1 is hypothesized to influence plant reproduction (Grover *et al*., 2020; Long *et al*., 2021), and it is also possible that the CLV1 and NRPD1 pathways intersect via siRNA movement.

The *brclv1-1* is an allele causing multilocular fruit production. Its potential of inducing seed yield could be maximized if the second site suppressor which reduces the internal gynoecia was found. Given the strong phenotype, it also provides fascinating molecular and genetic resources for resolving how CLV1 is involved in carpel development and regulation.

## Materials and Methods

### Plant materials and growth condition

All plants were grown in a greenhouse (*B. rapa*) or growth chamber (*Arabidopsis*) at 18° C with 16 hr light. Unless mentioned otherwise, all *B. rapa* plants used throughout the study are in the R-o-18 background. *clv11-11, clv1-12*, and *clv1-13* are null *clv1* mutant characterized before (Diévart *et al*., 2003). *clv1-12* and *clv1-13* were obtained from ABRC. *clv1-11* was a gift from Professor Zachary Nimchuk.

### Ethyl methanesulfonate (EMS) mutagenesis and isolation of *pomona*

EMS treatment of *nrpd1a-2* seed was performed as described in Stephenson *et al*., 2010. Briefly, *nrpd1a-2* seeds (4-6 generations backcrossed from the original TILLING mutation) were treated with 0.2% EMS for 16 hours, followed by multiple washing with 0.02% Tween20. The treated M1 seeds were grown and selfed seed were collected. Pools of M2 seed were propagated and M2 individuals with increased seed production were selected. *Brclv1* was recovered in this M2 generation and characterized in multiple M3 individuals.

### Map-based cloning of multilocular phenotype

M3 *brclv1* flowers were emasculated 1 day prior to anthesis, and then were pollinated with R500 pollen. The resulting F1 individuals were self-pollinated to generate an F2 population. F2 individuals from a single F1 parent were sown and phenotyped for locule number.

DNA was extracted from 46 multilocular F2 individuals using GeneJET Plant Genomic DNA Purification kit (Thermo Scientific). After quantification by Nanodrop, equal amounts of DNA were pooled and 2 µg of pooled DNA in 100 µL was fragmented by Bioruptor Pico (30s ON /90s OFF, 3 cycles, centrifuge 10s; repeat 1 time). 40 µl of the fragmented DNA was used for library construction with a NEXTflex PCR-Free DNA-Seq Kit (Bioo Scientific) according to the manufacturer instructions. Barcoded libraries were quantified using a Bioanalyzer (Agilent Technologies), and paired-end sequenced on an Illumina NextSeq 500 at The University of Arizona Genetics Core.

### Sequencing read and variant analyses

Trimmed reads were first aligned to the *B. rapa* R500 genome (version 1.2; Greenham et al., 2020) using Bowtie v2.2.4 (Langmead and Salzberg, 2012) and SNPs were used to identify linkage with the causal mutation. Variant calling was conducted by bcftools mpileup and only SNPs with quality score ≥ 30 and read depth > 10 were retained. bcftools query was used to extract information of each variant call and the results were further parsed using R to compute allele frequencies. For markers ≥ 50.0 map units from the causal mutation, the allele frequencies were expected to be ∼ 0.5; while markers linked to the mutation have higher allele frequencies. This analysis demonstrated linkage to the lower arm of chromosome 7.

To identify EMS-induced mutations in this region, read alignment and variant calling were conducted as described above but using the *B. rapa* R-o-18 genome (version 2.3). Only G-to-A and C-to-T, and mutant allele frequency > 0.9 SNPs within the mapping interval were retained for further analysis. To predict the effects of mutations, SnpRff was used to annotate the variants and predict their effects on genes (Cingolani *et al*., 2012).

### Genotyping *BrCLV3, BrCLV1*, and *BrCPEPM*

Genotyping primers were designed by dCAPS Finder 2.0 http://helix.wustl.edu/dcaps/ (Neff et al., 2002). The R-o-18 and R500 alleles at *BrCLV3* were genotyped with GATCGGAATCGGGAAGATGACAA and CGACGCTGATGAGGATCAACGC and digested with *HindIII. brclv1-1* was genotyped with GATCGGAATCGGGAAGATGACAA and CGACGCTGATGAGGATCAACGC and digested with *AluI*. Additional alleles at *BrCLV1* from the RevGen UK TILLING population (Stephenson *et al*., 2010) were verified through amplifying and sequencing regions of *BrCLV1* (forward1: AAACATCCCACCAGAACTCTCC and reverse1: GGAGATTAAGGAAGTGCAGCGAGA or forward2: gCTAACAATTGGTTTACCGGTTTAAGC and reverse2: GCTATCGAAGACATACACTCAGAAGCA). *BrCPEPM* was genotyped with GCACAAAGGGTTTCAGAGTCTGC and CTCCAAGTGACCTCTCGTGGATC and digested with *BamHI*.

### Cloning constructs and transformation

All cloning constructs were based on a pGGZ003 vector containing a UBQ10 promoter and a UBQ terminator (Lampropoulos *et al*., 2013). To remove the existing coding sequence between the promoter and terminator, the vector was digested with *NotI* and *BamHI* and gel purified. Gibson assembly modules were created via PCR amplification from genomic DNA of a 4005-bp *AtProCLV1* and approximately 3.3 kb sequences from *AtCLV1, BrCLV1*, or *brclv1-1* using primers in **Table S2**. The modules were joined to the backbone through a Gibson assembly reaction at 50°C for 1.5 hr and immediately transformed into *E. coli*. Sequences were confirmed by enzyme digestion and sequencing before transformation into chemically competent *Agrobacterium tumefaciens* GV3101 carrying the pSOUP helper plasmid for transformation of Arabidopsis by floral dip (Clough and Bent, 1998). T1 seeds were selected based on Basta resistance, and T2 plants from lines with a single insertion site were used for phenotyping.

### Yield assessment

The number of seeds per silique were counted from siliques collected from the main stem. The total number of seeds was assayed from all seeds collected from the whole plant.

### Evolutionary comparison

The genomic sequence of genes in the mapping interval were used as query sequences to search against the Arabidopsis (TAIR 10) genome using BLAST. The best hit with e-value lower than 1e-10 was reported. Retention of homoeologous copies among three subgenomes of *B. rapa* was retrieved from the BRAD database (http://brassicadb.cn/).

CLV1 orthologs in other species were obtained from Phytozome v13 (https://phytozome-next.jgi.doe.gov) (Goodstein *et al*., 2012) with following PAC gene identifiers: *C. rubella*, 20904558; *C. sativa*, 16979492, *E. salsugineum*, 20191806; *C. papaya*, 16415260; *Camellia sinensis*, 18092691; *R. communis*, 16822209: *C. esculenta*, 32335358; *M. truncatula*, 31112997; *F. vesca*, 27262373; *P. persica*, 32118556; *E. grandis*, 32053920; *A. hypochondriacus*, 32828513; *A. coerulea*, 33062625; and *S. polyrhiza*, 31521071.

### Accession number

Sequence data from this article can be found in NCBI SRA under accession number PRJNA882476.

## Acknowledgments

This work is supported by project number ARZT-3039860-G25-578 from the USDA National Institute of Food and Agriculture to RAM. We are grateful to Keign Vedvick for assistance with the EMS mutant screens, and to Renee Grambihler, Collin Eckhauser, and Jeff Clark for assisting with phenotyping and seed counting. Thanks also go to Dr. Frans Tax for reviewing the manuscript, Dr. Michael Ottman for use of his seed counter, and to Dr. Zachary Nimchuk for the *atclv1-11 line*. DNA sequencing was conducted by the University of Arizona Genetics Core.

## Notes

### Competing Interest Statement

The authors have declared no competing interest.

