## Supplemental Figure for "A novel CLAVATA1 mutation causes multilocularity in *Brassica rapa*"

### Supplemental Materials for Chow H, *et al.* 2022

**Contents:**

Supplemental Tables 1-2

Supplemental Figures 1-5

**Supplemental Table 1. Sequencing statistics.**

| | |
|---|---|
| Pair-end reads | 207,047,206 |
| Pair-end read QC passed | 100% |
| Mapped reads | 201,084,246 |
| Alignment rate (%) | 97.10% |
| SNPs identified | 279,071 |

**Supplemental Table 2. Primers for cloning.**

| Name | Primer sequence (5' to 3') |
|---|---|
| AtPro-F[#&] | GCGGCCGCGGGGTTTATCTGAATTGGATGGTG |
| AtPro-R[&] | AGAAGGTGAGTTTTCAGAAGTCTCATTTTTTTAGTGTCCTCTCAGTGAGAAAG |
| BrCLV1-F[&] | CTTTCTCACTGAGAGGACACTAAAAAAATGAGACTTCTGAAAACTCACCTTCT |
| BrCLV1-R[&] | ACAGGGAATGAAGGTAAAGGTTATCTAGGAAGTTTAAAAGCCATCATCAATAAGCAT |
| AtCLV1-R[#] | CAATTTACAGGGAATGAAGGTAAAGGTTATCTAGCCCTAATTCACATATTCTAAACAGTCTAGAC |

[#] denote primers used for *AtProCLV1-AtCLV1* cloning while [&] are used for *AtProCLV1-BrCLV1* cloning.

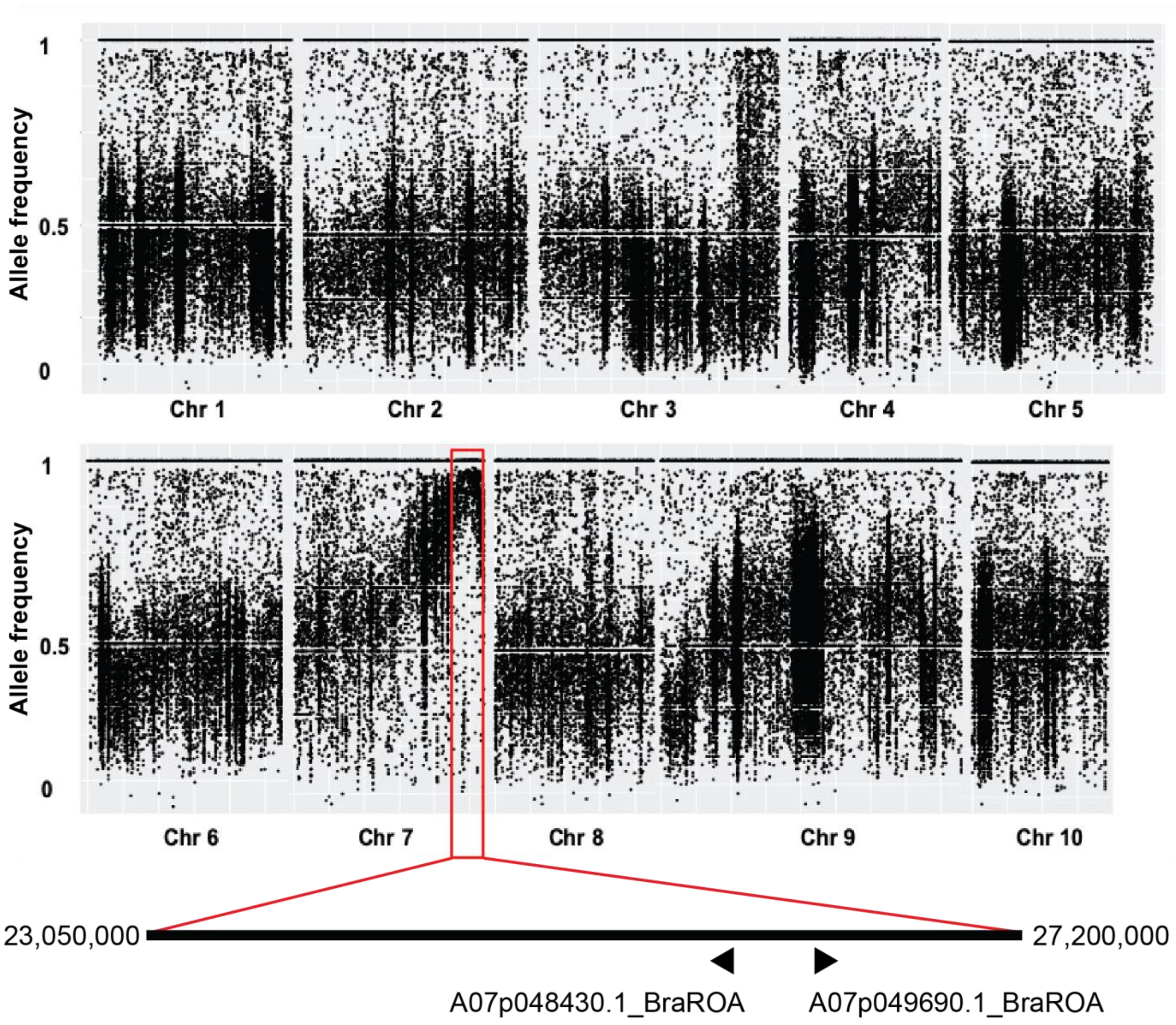


**Supplemental Figure 1. *A07p048430.1_BraROA* is one of the top candidates in the mapping interval.**

(A) Allele frequencies of all SNP variants over *B. rapa* chromosomes among pooled multilocular individuals from an F2 population. The peak (red box) indicates that *POMONA* is on chromosome 7.

(B) Schematic representation of the mapping interval and arrows represent the position of the two top candidate genes.

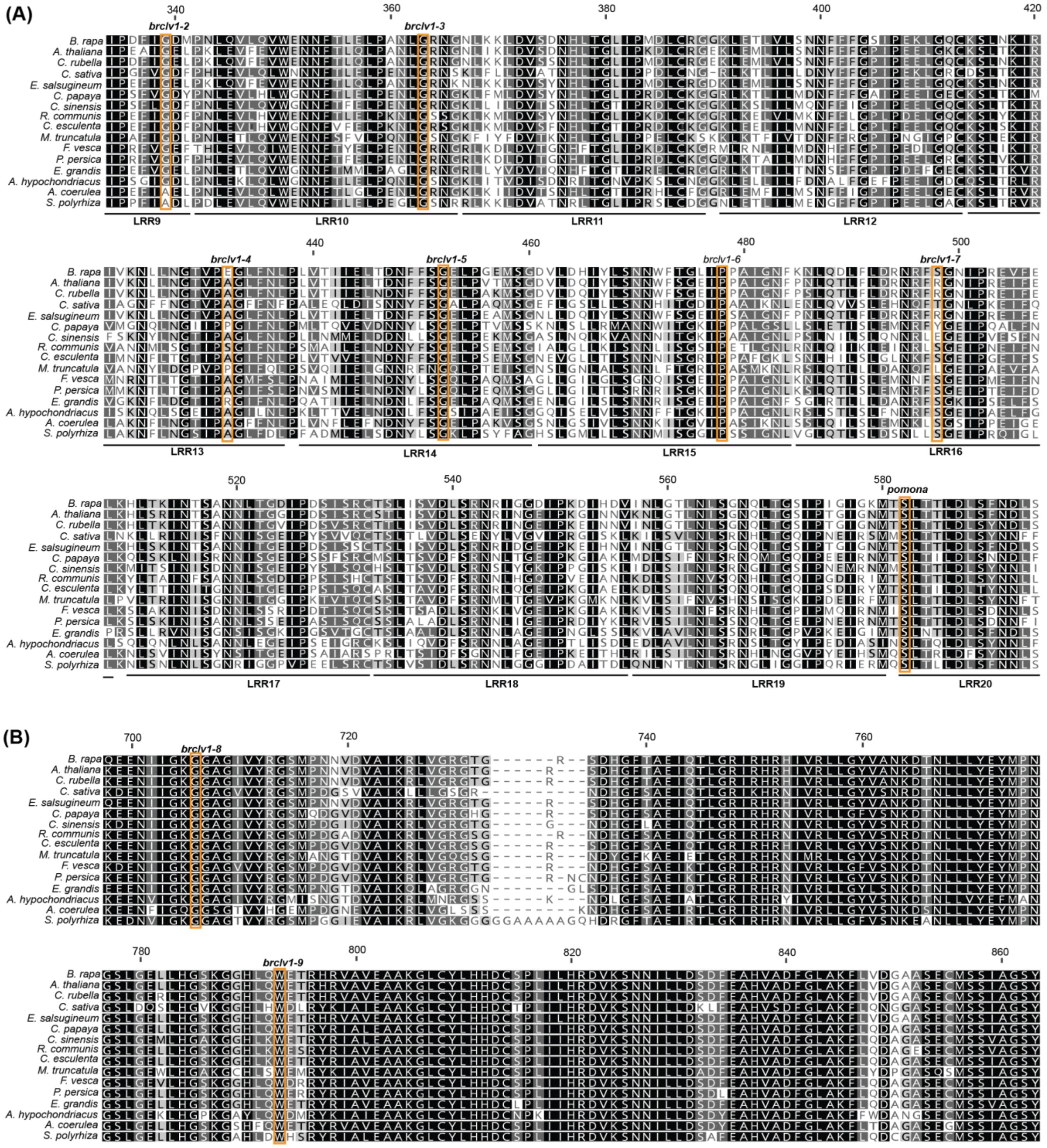


**Supplemental Figure 2. Amino acid alignment of BrCLV1 with orthologs in the eudicots. Comparison of amino acid sequences of (A) LRR domain (position 334 – 589) and (B) kinase domain (position 684 – 863) in** *Brassica rapa* (*B. rapa*), *Arabidopsis thaliana* (*A. thaliana*), *Capsella rubella* (*C. rubella*), *Camelina sativa* (*C. sativa*), *Eutrema salsugineum* (*E. salsugineum*), *Carica papaya* (*C. papaya*), *Camellia sinensis* (*C. sinensis*), *Ricinus communis* (*R. communis*), *Colocasia esculenta* (*C. esculenta*), *Medicago truncatula* (*M. truncatula*), *Fragaria vesca* (*F. vesca*), *Prunus persica* (*P. persica*), *Eucalyptus grandis* (*E. grandis*), *Amaranthus hypochondriacus* (*A. hypochondriacus*), *Aquilegia coerulea* (*A. coerulea*), and *Spirodela polyrhiza* (*S. polyrhiza*). Identical amino acid residues are rendered black. Similar residues in at least 80% all comparing species are highlighted in dark-grey, whereas similar residues in at least 60% but lower than 80% similarity are highlighted in light-grey. Mutated residues used in the experiment are indicated above the alignment. Number on the top refers to the position of BrCLV1 amino acid position.

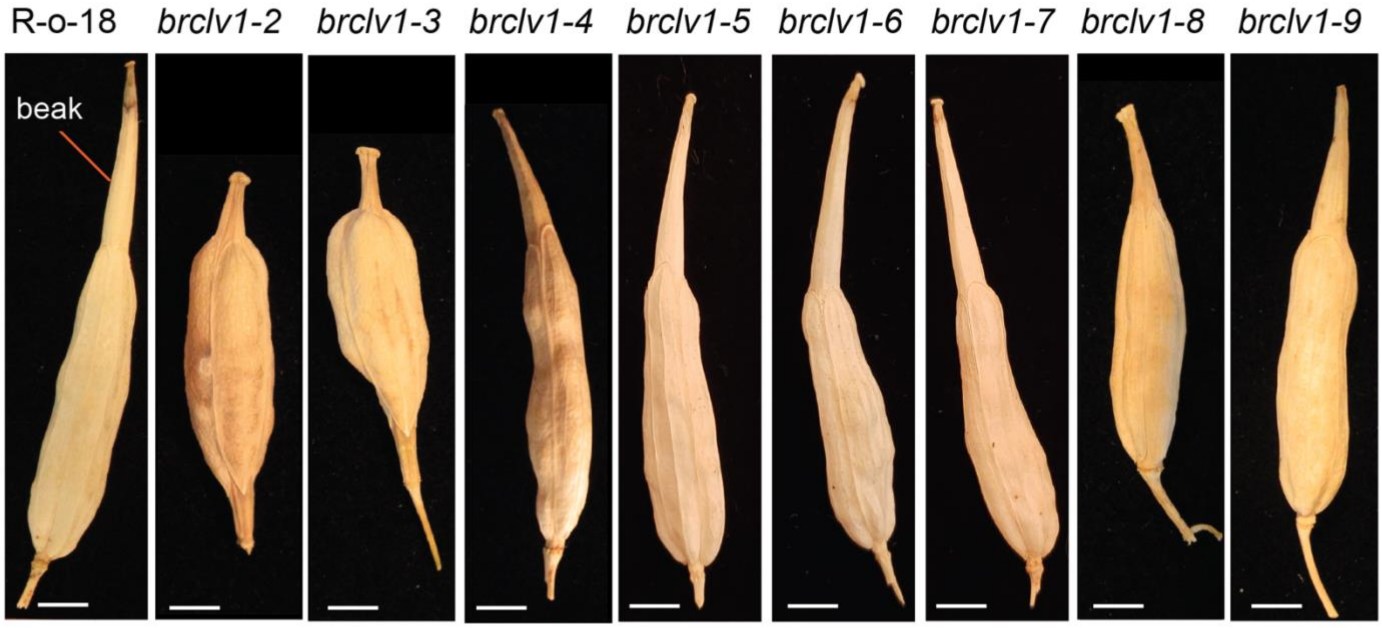


**Supplemental Figure 3. *Brclv1-2*, *Brclv1-3*, and *Brclv1-8* cause changes in silique phenotype.**
Representative of siliques from left to right: R-o-18 (WT), *brclv1-2*, *brclv1-3*, *brclv1-4, brclv1-5*, *brclv1-6, brclv1-7*, *brclv1-8, and brclv1-9.* Scale bar = 0.5 cm.

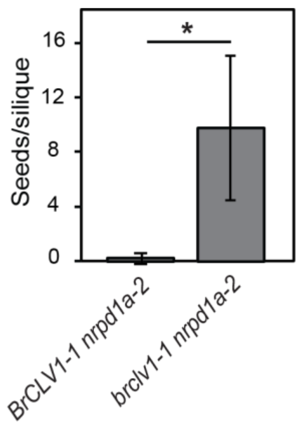


**Supplemental Figure 4. *brclv1-1* causes more seed yield in the *nrpd1a-2*.**
Boxplot of seeds/silique number per plant in the *brclv1-1/nrpd1a-2* and *BRCLV1-1/nrpd1a-2*. Asterisk above denotes significant difference according to two-sample t-test (p-value 5.99e-11< 0.05).

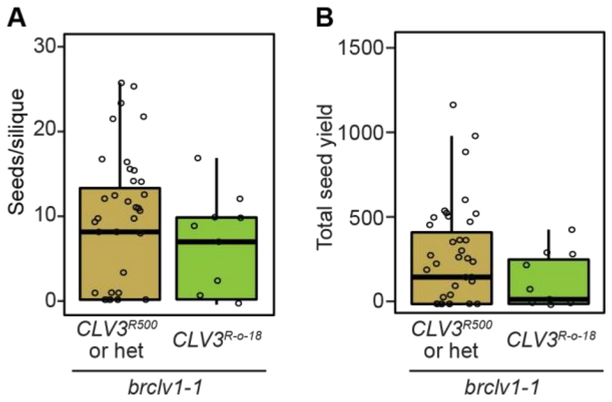


**Supplemental Figure 5. BrCLV1 and BrCLV3 do not act synergistically in boosting seed set.** No significant difference between comparing groups according to two-sample t-test.
